# One-Tip enables comprehensive proteome coverage in minimal cells and single zygotes

**DOI:** 10.1101/2023.08.10.552756

**Authors:** Zilu Ye, Pierre Sabatier, Javier Martin-Gonzalez, Akihiro Eguchi, Dorte B. Bekker-Jensen, Nicolai Bache, Jesper V. Olsen

**Author notes:** These authors contributed equally (Zilu Ye, Pierre Sabatier).

## Abstract

We present One-Tip, a lossless proteomics methodology that seamlessly combines swift, one-pot sample preparation with narrow-window data-independent acquisition mass spectrometric analysis. With simplest sample processing, One-Tip reproducibly identifies > 9,000 proteins from ∼1000 cells and ∼ 6,000 proteins in a single mouse zygote with a throughput of 40 samples-per-day. This easy-to-use method expands capabilities of proteomics research by enabling greater depth, scalability and throughput covering low to high input samples.

## Main

Conventional bottom-up proteomics workflows typically involve multi-step processes, incorporating cell lysis and protein extraction from millions of cells, reduction and alkylation of cysteines, protein digestion, sample cleanup and SpeedVac, followed by optional offline peptide fractionation prior to online liquid chromatography tandem mass spectrometry (LC-MS/MS) acquisition^1^. The different steps introduce several challenges, including potential sample contamination and loss leading to reproducibility problems, considerable time and cost investments, and the necessity for specialized expertise and training. Consequentially, these barriers often discourage biomedical scientists from undertaking MS-based proteomics experiments. While certain one-pot workflows such as iST^2^ have been introduced to simplify sample preparation for bulk proteomics—akin to single-cell proteomics^3^—these methods still require substantial sample input, separated cell lysis, multiple liquid handling steps, sample transfer, and prolonged LC-MS/MS analysis time^2, 4, 5^. These factors result in limited throughput, reproducibility, applicability and proteome depth.

In this study, we developed One-Tip, a streamlined workflow designed to circumvent these issues. The One-Tip methodology only requires two pipetting steps, utilizing a commercially available Evotip™: one for the lysis and digestion buffer, and the other for the cell suspension in PBS (**Fig. 1a**). After a mere hour-long incubation in a water-filled Evotip box and standard Evotip preparation, the One-Tip samples are ready for LC-MS/MS analysis. The combined lysis and trypsin digestion master mix buffer includes an MS-compatible surfactant, n-Dodecyl-β-D-Maltoside (DDM), which is a water soluble non-ionic detergent effective in cell membrane lysis and solubilization of proteins without denaturation.

**Figure 1.**
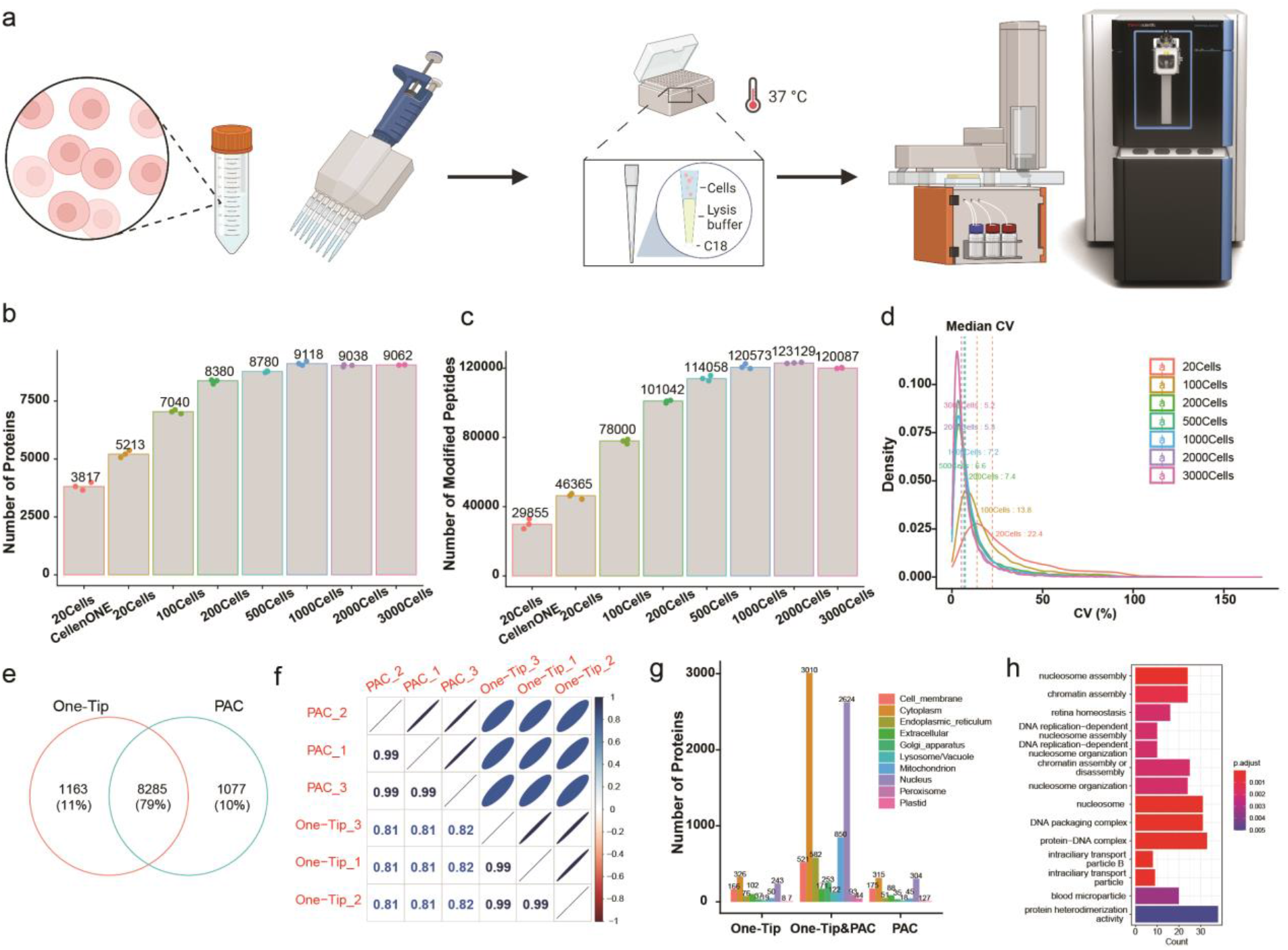
One-tip achieves near-complete proteome depth in single-shot analysis. **a)** Graphic depiction of One-Tip workflow. **b-c**) Number of proteins (**b**) and peptides (**c**) identified in different number of cells. Digestion time was 2h in these samples. **d**) Distribution of coefficient of variance between triplicates in different number of cells. **e**) Overlap of proteins identified in One-Tip (1000 cells) and PAC (1000ng) workflows. **f**) Correlation of protein abundances between One-Tip and PAC samples. **g**) Subcellular localization of proteins identified in One-Tip and PAC samples. **h**) GO over-representation enrichment of proteins identified only in One-Tip samples.

Conceptually, One-Tip represents the simplest workflow for proteomics, as it necessitates no further sample handling and directly integrates sample preparation with the LC-MS analysis without any sample losses and variation. Low cell input numbers result in high proteome coverage and circumvents the need for sonication and additional buffer exchanges. This results in a total sample preparation time of approximately 70 minutes including a digestion time of 60 minutes. Moreover, the One-Tip workflow is not limited in throughput; even manual handling can efficiently prepare thousands of samples daily by using multi-channel pipetting and it is also easily automated on liquid handling robots. Lastly, we couple the One-Tip workflow with a finely-tuned LC-MS/MS system to guarantee superior analytical performance. This is based on the highly sensitive Whisper LC method on the EvoSep One LC and our newly developed narrow-window Data Independent Acquisition (DIA) method on an Orbitrap Astral mass spectrometer^6^.

To test sensitivity and reproducibility of the One-Tip methodology, we initially deployed it with increasing numbers of HeLa cells, analyzing 20, 100, 200, 500, 1,000, 2,000, and 3,000 cells, respectively. Note, these cell quantities were based on cell counter estimations, and therefore are approximations. One-Tip exhibited an impressive proteome depth for single-shot analysis in half-an-hour LC-MS/MS runs, identifying over 5,000 protein groups (hereafter referred to as proteins) and over 46,000 modified peptides (hereafter referred to as peptides) from approximately 20 cells (**Fig. 1b**). This performance surpasses the proteome coverage currently achieved by the isolation of precisely 20 cells using the CellenONE, a dedicated single-cell proteomics preparation system, which is likely related to sample losses in a 96 well plate and transfer to the LC-MS system. Furthermore, a starting quantity of ∼1,000 cells nearly covered the complete proteome, with over 9,000 proteins and 120,000 peptides identified (**Fig. 1b, 1c**). We subsequently evaluated the trypsin-digestion efficiency by varying the digestion time from 1 to 4 hours. Even a digestion time as short as 1 hour was sufficient to effectively digest the entire proteome, resulting in high sequence coverage with a missed cleavage rate of less than 25% in 20 cells and approximately 30% for larger cell quantities (**Supplementary Figure 1-3**). One-Tip also demonstrated high reproducibility and quantification precision, with a remarkable coefficient of variance of 5-8% and a Pearson correlation exceeding 0.99 for proteins in samples more than 100 cells (**Fig. 1d, Supplementary Figure 4**).

Despite its simple and swift workflow, the analytical performance of One-Tip with one thousand cells is comparable to that of bulk proteomics samples prepared using a more sophisticated method, Protein Aggregation Capture (PAC)^7^, when combined with our narrow-window DIA method on the Orbitrap Astral mass spectrometer. Compared to 1,000-ng PAC-digested HeLa lysate analyzed by LC-MS/MS in >1 hour, the One-Tip method applied to 1,000-cell samples in a half-an-hour LC-MS/MS run identified a similar number of proteins with substantial overlap between them and very high quantification precision (**Fig. 1e, 1f**). Interestingly, both the PAC and One-Tip methods identified slightly more than 1,000 unique proteins, but exhibiting no considerable bias towards specific subcellular localizations (**Fig. 1g**), and slight preference for particular biological functions such as nucleosome assembly in proteins identified only in One-Pot samples(**Fig. 1h**).

To highlight the versatility and sensitivity further of the One-tip workflow, we next studied the development of mouse pre-implantation embryos at single-cell level from oocyte and zygote to 4-cell stage. The zona pellucida, the extracellular glycoproteinaceous coat of oocytes, zygotes, 2-cell and 4-cell stages, was chemically removed and single cells were dissociated under a microscope and dispensed individually into Evotips, pre-loaded with lysis and digestion buffer (**Fig. 2a**). This whole procedure was performed in one day for each batch of samples including the LC-MS/MS analysis. In total, 6216 protein groups were identified with a median of 5428, 5417, 5259 and 4779, for each stage, respectively (**Fig. 2b**). In comparison to the very recent analysis of human pre-implantation embryos from Dang et al. ^8^ in which the authors employed a sophisticated sample preparation workflow and LC gradient 4 times longer than the one used in this study, we identify ∼2300 more proteins and ∼50,000 more peptides as well as more peptide per protein on average (5.3 against 14.6), per stage (**Supplementary Figure 5a, 5b**). Moreover, the variation in protein number, the overlap (**Fig. 2b**) and correlation of protein quantities between samples from a same condition was greatly improved (**Supplementary Figures 5c, 5d**), demonstrating that our One-Tip workflow is faster, more sensitive, more reproducible and has better quantification accuracy.

**Figure 2.**
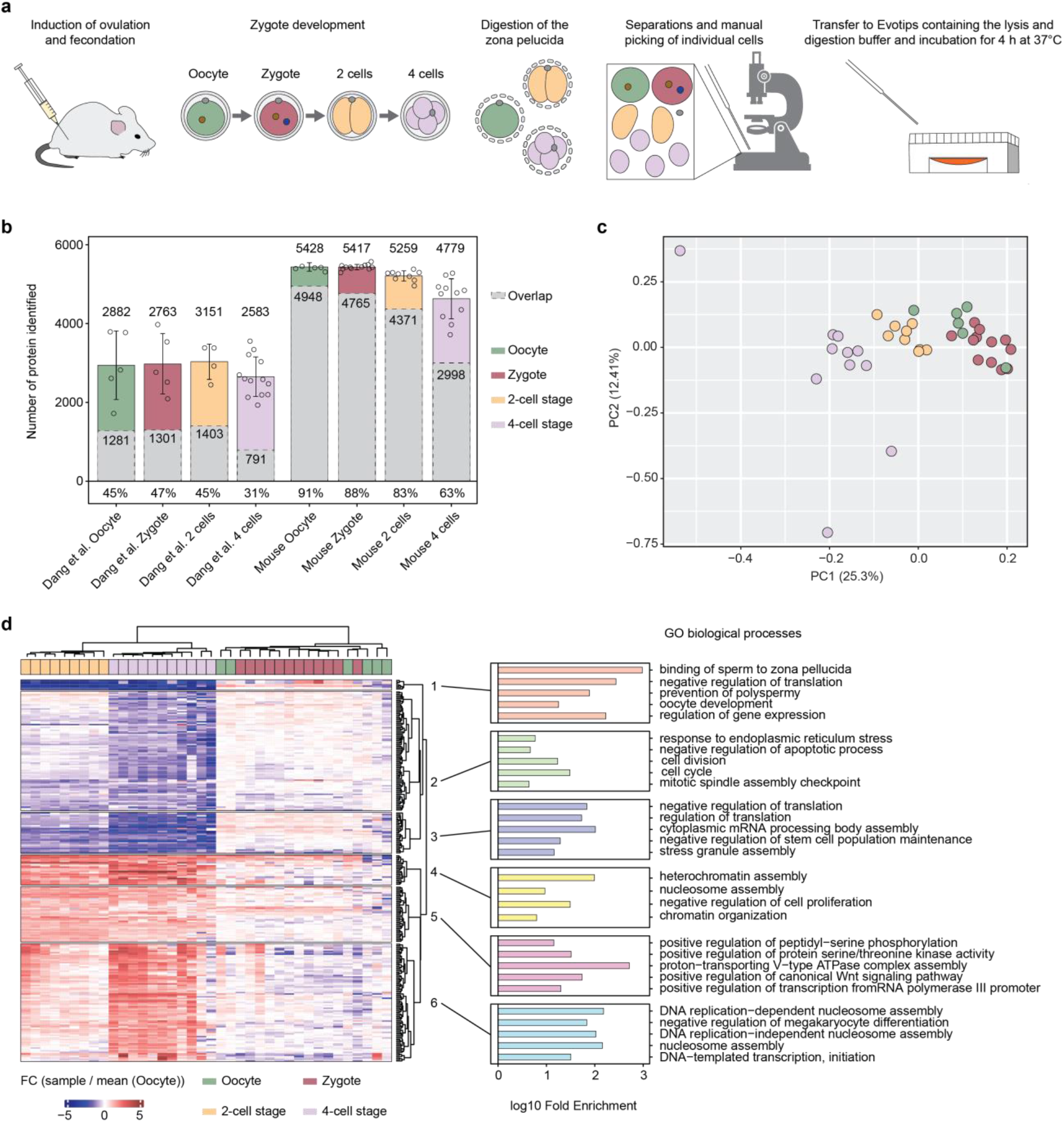
One-tip and LC-MS analysis of the mouse pre-implantation embryo until the 4-cell stage. (**a**) Workflow. (**b**) Number of proteins with quantified values in the study from Dang et al. and our study, and protein overlap between samples (in grey) from the same development stages. The mean number of proteins for each condition, the overlap and the mean percentage of the overlap compared to each sample are indicated. (**c**) PCA of the normalized protein intensities for each samples of our study. Only proteins that were quantified in all samples were considered (n = 2953). (**d**) Unsupervised hierarchical clustering using canberra and ward.D2 methods of proteins that shows significant regulation (p<0.05 from ANNOVA) and FC > 2 fold in any of the developmental stages compared to the oocyte group (left). GO biological processes enrichment analysis of the protein clusters from the heatmap (right). n=6 for oocyte, n=12 for zygote, n=9 for 2-cell stage and n=11 for 4-cell stage.

Finally, we investigated whether biologically meaningful information can be retrieved from the mouse embryo cell analysis. The principal component analysis (PCA) of normalized protein intensities showed that oocyte and zygote group together and the 2-cell and 4-cell stages are separated in a time-wise manner on PC1 (**Fig. 2c**). Interestingly, the summed protein intensities in each sample were equal between oocytes and zygotes while they were half those intensities in cells from the 2-cell stage and one fourth in cells from the 4-cell stage, matching the reduction in cell volume during the two successive cleavages of the zygote (**Supplementary Figure 5e**). We then performed unsupervised hierarchical clustering of differentially-expressed proteins that were up or downregulated in any of the conditions compared to oocytes (**Fig. 2d**). Each sample group was separated on the heatmap apart from the zygotes and oocytes as on the PCA. GO enrichment analysis of clusters highlights downregulation of proteins related to sperm binding to the zona pellucida, prevention of polyspermy, response to ER stress and upregulation of proteins related to DNA replication and heterochromatin assembly in 2-cell and 4-cell stages compared to oocytes, among other pathways. These results were globally in agreement with Dang et al. validating the quality of our analysis (**Fig. 2d**).

Taken together, our results demonstrate that the analytical depth achieved with the One-tip workflow is superior to standard workflows in the field while its simplicity, ease-of-use and scalability make it suitable for many experimental designs abrogating the need for sophisticated methods, specific equipment and expertise in proteomics during sample preparation. This workflow will greatly contribute to bridge the gap between proteomics and biology by making sample preparation for LC-MS analyses more streamlined, user-friendly, compatible with other platforms such as CellenONE and readily accessible to most biological applications.

## Online Methods

### HeLa Cell lines

HeLa cells were cultured in DMEM (Gibco, Invitrogen), supplemented with 10% fetal bovine serum, 100U/ml penicillin (Invitrogen), 100 μg/ml streptomycin (Invitrogen), at 37 °C, in a humidified incubator with 5% CO_2_. At ∼80% confluence. cells were detached using trypsin and washed twice with Phosphate Buffered Saline (PBS) from Gibco (Life Technologies), before being resuspended in degassed PBS.

### Detailed One-Pot sample preparation workflow

1. Determine HeLa cell concentration using a cell counter and dilute to the desired concentrations with PBS.
2. Prepare the Evotips following the vendor’s instructions:
  - Rinse: Wash dry Evotips with 20 μl of Solvent B (0.1% FA in acetonitrile) and centrifuge at 800 g for 60 seconds.
  - Condition: Soak the Evotips in 100 μl of 1-propanol until they turn pale white.
  - Equilibrate: Saturate the conditioned Evotips with 20 μl of Solvent A (0.1% FA in water) and centrifuge at 800 g for 60 seconds.
3. Pipette 5 μl of lysis and digestion buffer into the Evotips. The buffer contains 0.2% n-Dodecyl-β-D-Maltoside (DDM), 100 mM TEAB, 20 ng/μl Trypsin, and 10 ng/μl Lys-C.
4. Pipette 5 μl of cells into the Evotips.
5. Briefly centrifuge the Evotips at 50 g to mix the buffer and cells and prevent the formation of air bubbles.
6. Add water to the Evotip box to the level of the C18 resin in the Evotips.
7. Incubate the Evotip box at 37°C for 1 to 4 hours.
8. Continue with the vendor’s instructions with a slight modification:
  - Load: add 50 μl of Solvent A to the Evotips and centrifuge for 60 seconds at 800 g.
  - Wash: Rinse the tips with 20 μl of Solvent A and centrifuge for 60 seconds at 800 g.
  - Wet: Add 100 μl of Solvent A to the tips and centrifuge for 10 seconds at 800 g to keep the tips wet.

### HeLa cell isolation with CellenONE

HeLa cells were diluted with degassed PBS to ∼200 cells/μl. 20 cells were isolated into a 96 well-plate preloaded with 4 μl PBS using the CellenONE system. After isolation, samples were prepared following the One-Tip workflow.

### Isolation of mouse oocytes, zygotes, 2-cell and 4-cell stages

Animal work was conducted according to license no. 2021-15-0201-00851, approved by the Danish National Animal Experiments Inspectorate, and performed according to national and local guidelines. Mice were kept in designated rooms at a temperature of 22 °C (±2 °C), with a humidity of 55% (±10%), according to Danish regulations.

Ovulation of prepubescent (4-week old) C57BL/6NRj females was induced by intraperitoneal (IP) injection of PMSG (HOR-272, Prospec), 5IU/female, followed by IP injection of hCG (Chorulon Vet, Pharmacy), 5IU/female, 47h later. After the second injection, females were set in breeding with C57BL/6NRj stud males. Next morning, the females were euthanized and oviducts were dissected to harvest the cumuli containing zygotes and unfertilized oocytes. Cumuli were disaggregated by 10-minute incubation in Hyaluronidase (H4272, Sigma-Aldrich). Zygotes were sorted out assessed by the presence of the second polar body and cultured in KSOM medium (MR-106-D, Merc Millipore). On the day of harvesting a group of both zygotes and oocytes were processed for analysis, while other zygotes were incubated in KSOM at 37 °C and 5% CO_2_ for 24h or 48h. At the different developmental stages, oocytes and embryos were incubated in Tyrode’s Acidic Solution (T1788, Sigma-Aldrich) for 5 to 15 seconds in order to remove the zona pellucida. Naked oocyes and zygotes were washed in PBS (20012-027, Life Technologies) and individually transferred to Evotips using a 100 μm diameter glass capillary. 2-cell and 4-cell embryos were further disaggregated by 5-minute incubation in Ca2+-and Mg2+-free KSOM and pippeting with a 50 μm diameter glass capillary in PBS afterwards; individual blastomeres were then transferred to Evotips. The Evotips were preloaded with 2 μl of lysis and digestion buffer and 2 μl of PBS. Incubation time was 4 hours.

### LC-MS/MS

LC-MS/MS analysis was performed on an Orbitrap Astral MS coupled to an EvoSep One system (EvoSep Biosystems). Samples were analyzed in 40SPD (31-min gradient) using a commercial analytical column (Aurora Elite TS, IonOpticks) interfaced online using an EASY-Spray™ source. The Orbitrap Astral MS was operated at a full MS resolution of 240,000 with a full scan range of 380 − 980 m/z when stated. The full MS AGC was set to 500%. MS/MS scans were recorded with 2Th isolation window, 3 ms maximum ion injection time (IIT) for HeLa samples and 4Th and 6ms IIT for mouse embryonic samples. MS/MS scanning range was from 380-980 m/z were used. The isolated ions were fragmented using HCD with 27% NCE.

### MS data analysis

Raw files were analyzed in Spectronaut v18 (Biognosys) with a library-free approach (directDIA+) using the human reference database (Uniprot 2022 release, 20,588 sequences) for HeLa samples and the mouse reference database (Uniprot 2022 release, 21,989 sequences) for the mouse embryo cells complemented with common contaminants (246 sequences). Note, cysteine carbamylation was not set as a modification, whereas methionine oxidation and protein N-termini acetylation were set as variable modifications. Precursor filtering was set as Q-value, cross run normalization was checked. Each experiment with different number of cells was analyzed separately, and samples from different digestion times were search with and without enabling method evaluation and indicating the different conditions (each one with n=3 experimental replicates) in the condition setup tab.

