## Supplementary Figures for "One-Tip enables comprehensive proteome coverage in minimal cells and single zygotes"

a

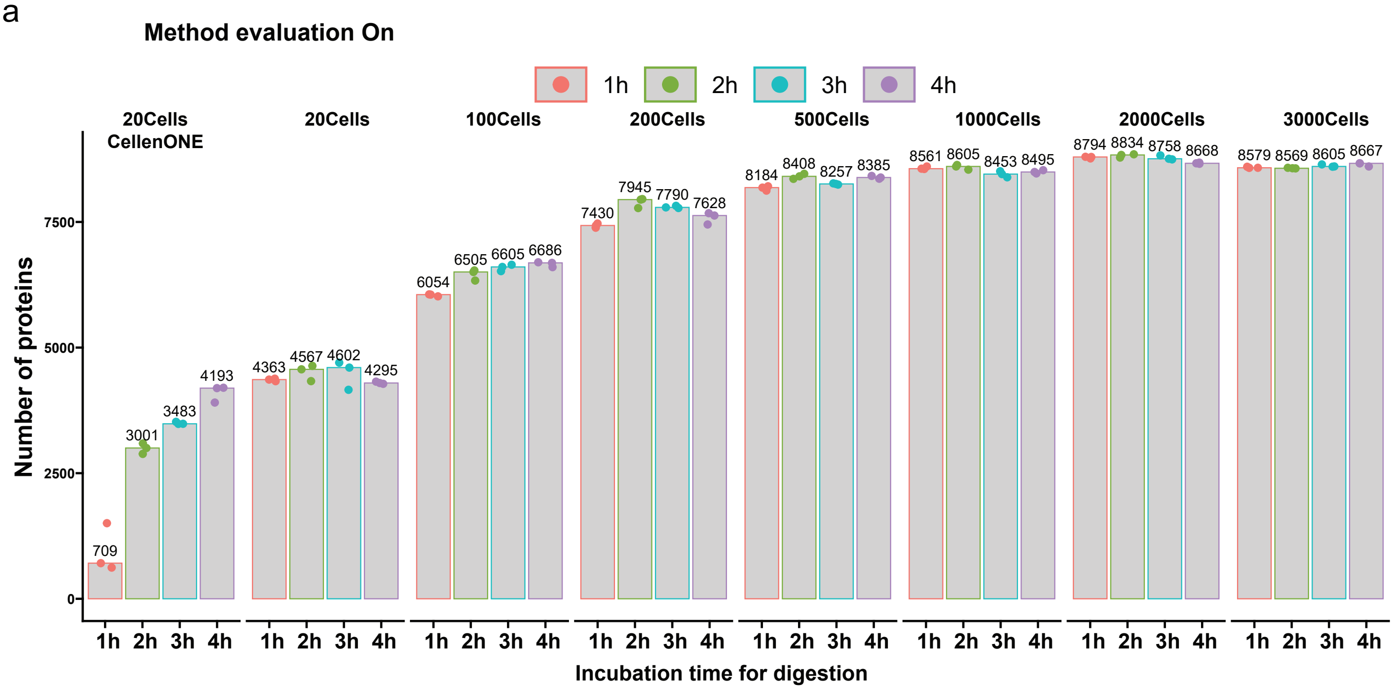


b

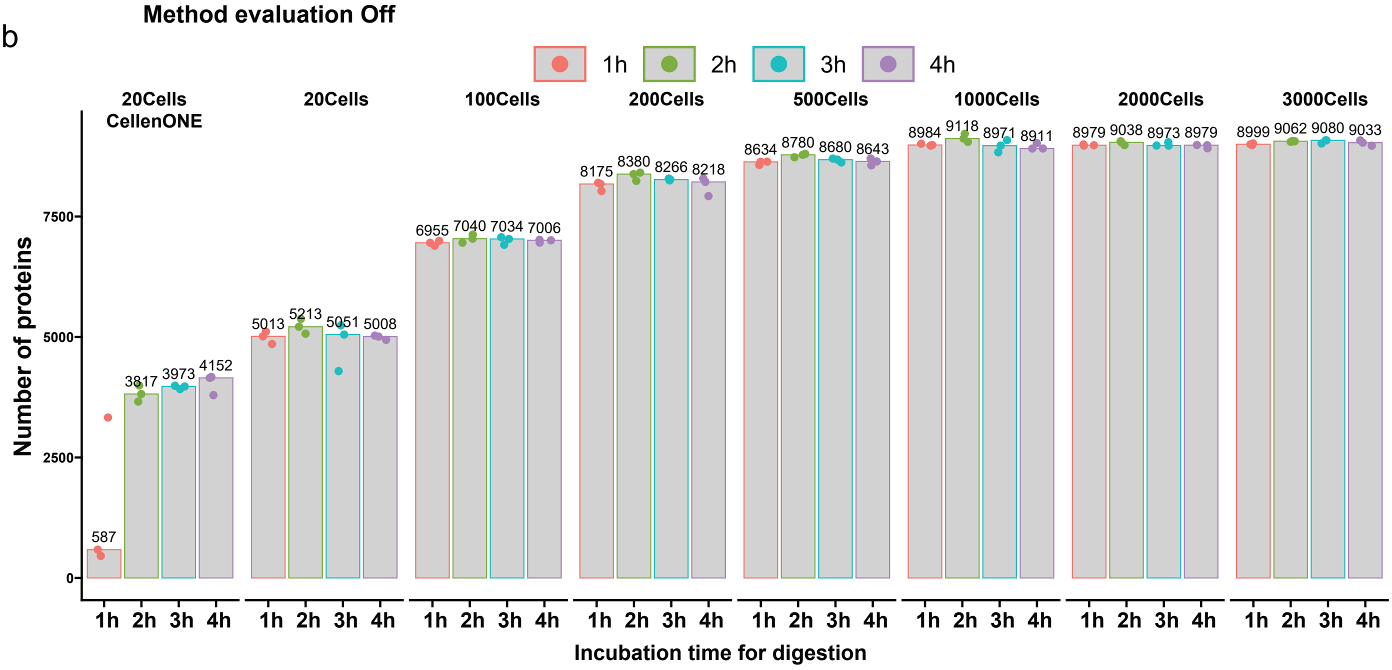


**Supplementary Figure 1**| Identification count of proteins across varying cell quantities using the One-Tip method. (**a**) With the 'Method Evaluation' option in Spectronaut enabled. (**b**) With the 'Method Evaluation' option in Spectronaut disabled. All samples underwent digestion times ranging from 1 to 4 hours.

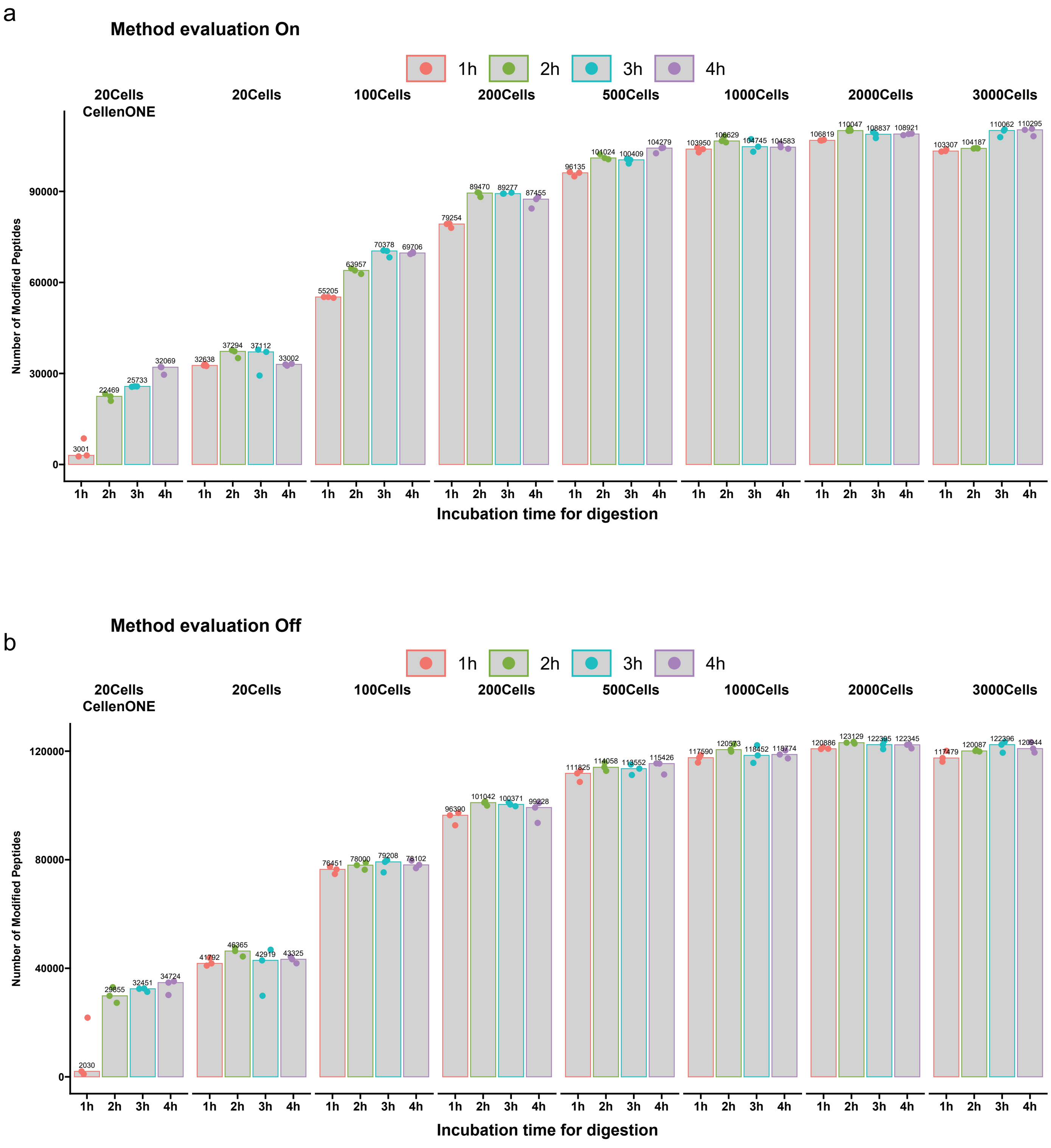


**Supplementary Figure 2** | Identification count of peptides across varying cell quantities using the One-Tip method. (**a**) With the 'Method Evaluation' option in Spectronaut enabled. (**b**) With the 'Method Evaluation' option in Spectronaut disabled. All samples underwent digestion times ranging from 1 to 4 hours.

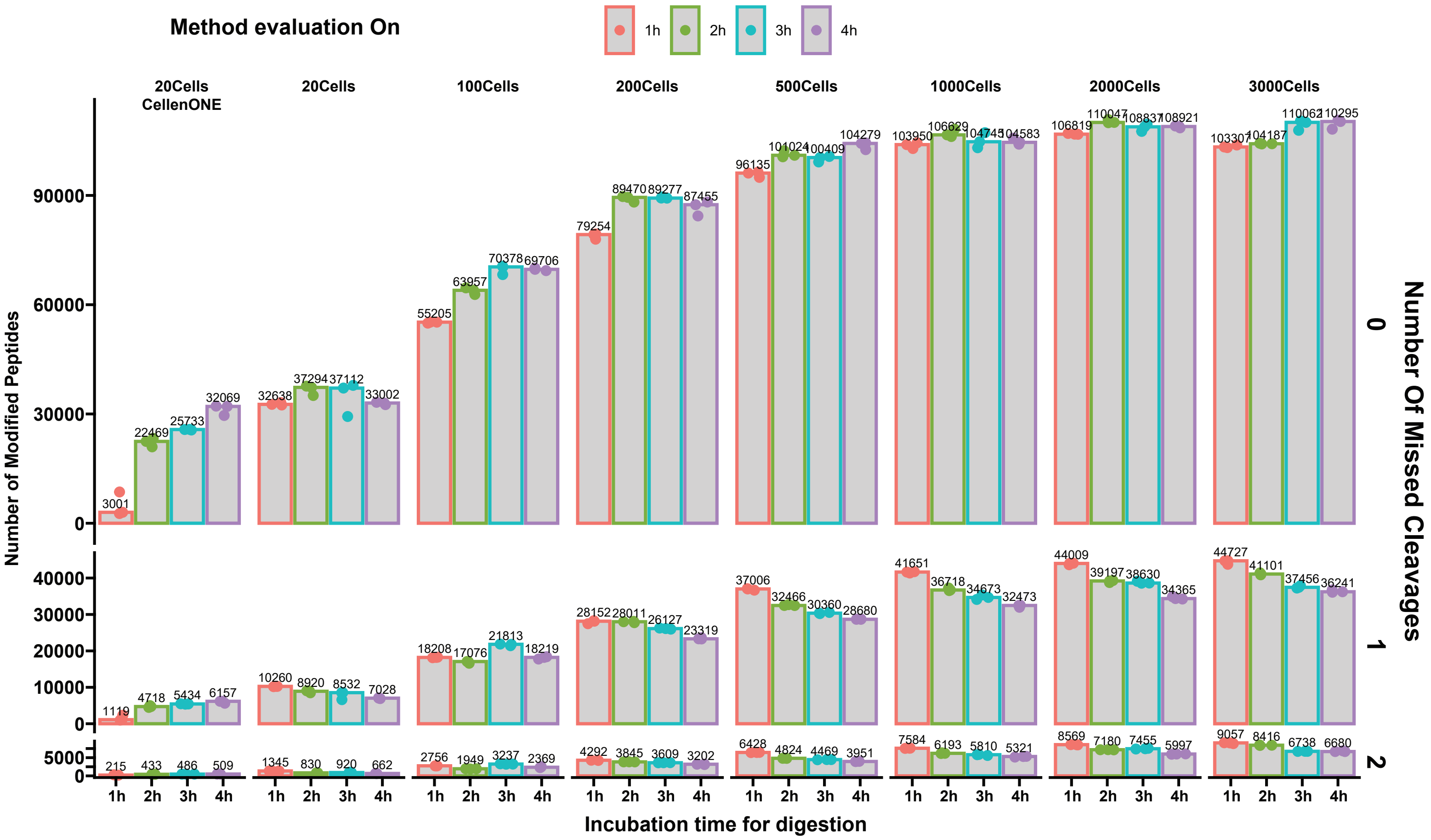


**Supplementary Figure 3** | Count of peptides with differing numbers of missed cleavages, analyzed with the 'Method Evaluation' option in Spectronaut enabled. The digestion duration varied from 1 to 4 hours.

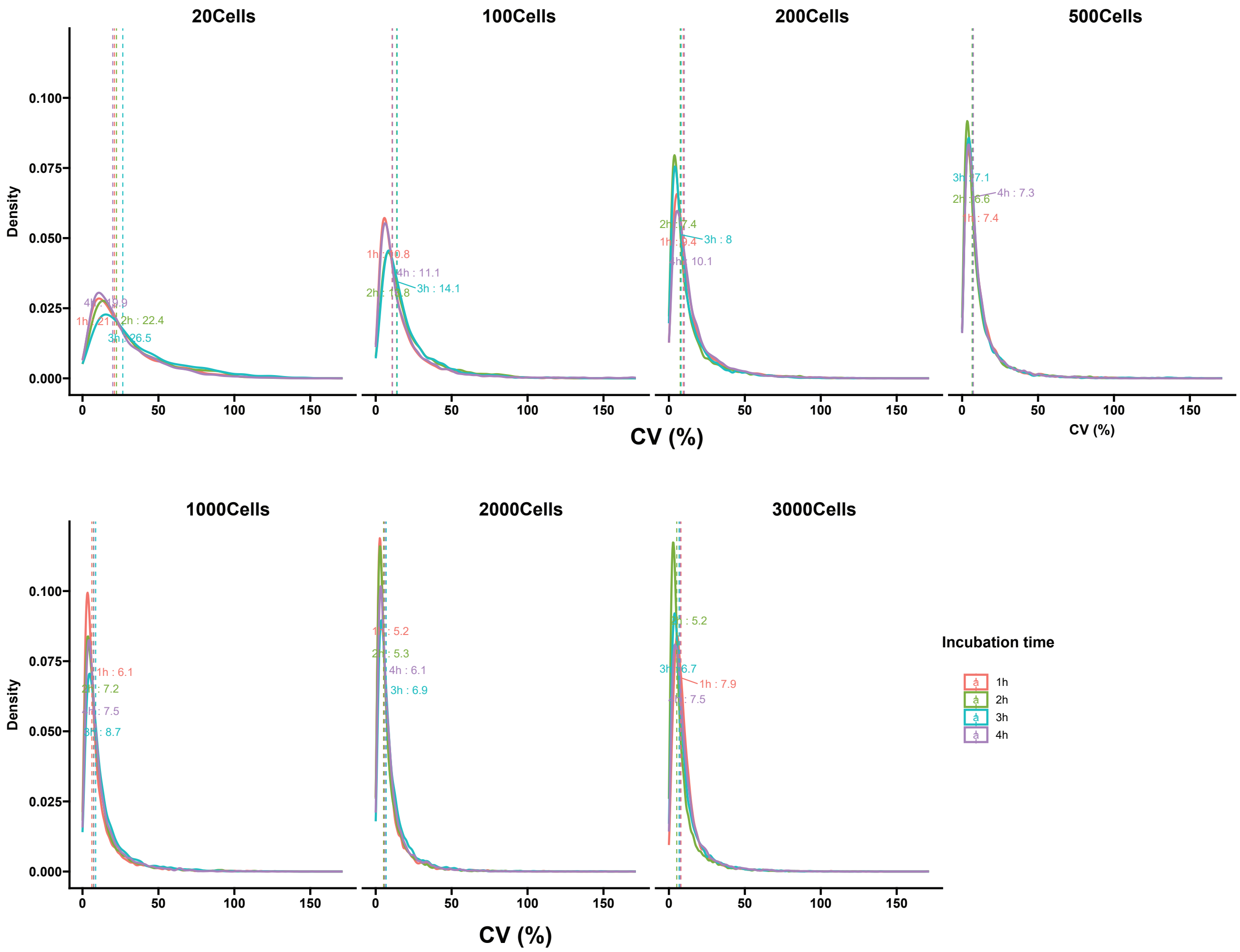


**Supplementary Figure 4** | Distribution of coefficients of variation (CV) across different cell quantities processed via the One-Tip method. Digestion periods spanned 1 to 4 hours, and median CV values are annotated.

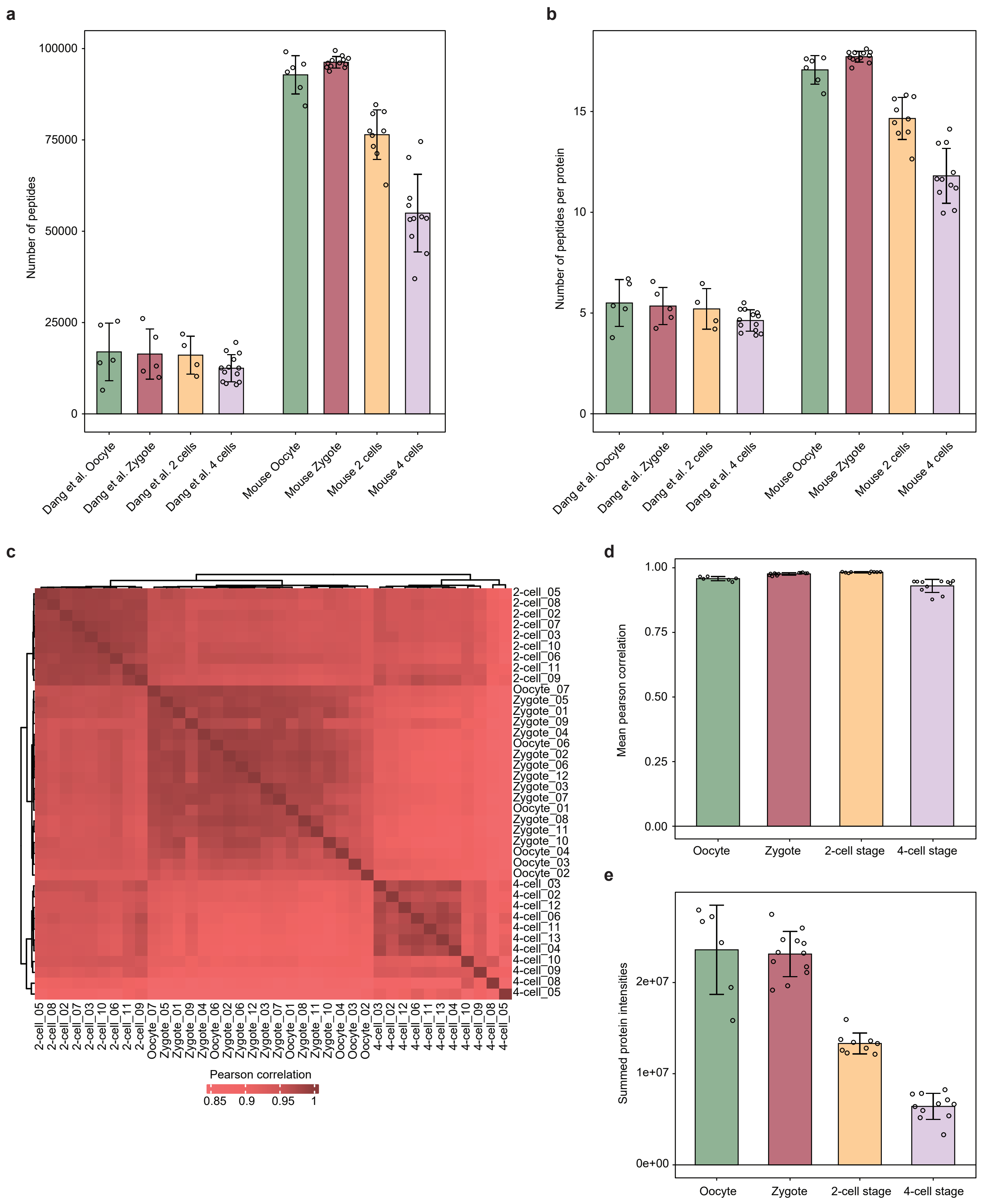


**Supplementary Figure 5 | Number of peptides, correlation of protein intensities between samples and summed protein intensities.** (**a**) Number of peptides per sample in Dang et al. study and our study. (**b**) Same with the number of peptides per protein. (**c**) Unsupervised hierarchical clustering using canberra and ward.D2 methods of the correlation of normalized protein abundances between each sample. (**d**) Mean correlation of normalized protein abundances between samples within each sample groups (oocyte, zygote, 2-cell stage, 4-cell stage). (**e**) Summed protein intensities for each group of samples. Error bars represent ± the standard deviation of the mean.
